# E2F transcription factors promote tumorigenicity in pancreatic ductal adenocarcinoma

**DOI:** 10.1101/2023.11.10.566445

**Authors:** Ludivine Bertonnier-Brouty, Jonas Andersson, Tuomas Kaprio, Jaana Hagström, Sara Bsharat, Olof Asplund, Gad Hatem, Caj Haglund, Hanna Seppänen, Rashmi B Prasad, Isabella Artner

## Abstract

Pancreatic ductal adenocarcinoma (PDAC) is one of the most lethal cancers with limited treatment options, illustrating an urgent need to identify new drugable targets in PDACs. Using the similarities between tumor development and normal embryonic development, which is accompanied by rapid cell expansion, we identified embryonic signalling pathways that were reinitiated during tumor formation and expansion. Here, we report that the transcription factors E2F1 and E2F8 are potential key regulators in PDAC. E2F1 and E2F8 RNA expression is mainly localized in proliferating cells in the developing pancreas and in malignant ductal cells in PDAC. Silencing of E2F1 and E2F8 in PANC-1 pancreatic tumor cells inhibited cell proliferation and impaired cell spreading and migration. Moreover, loss of E2F1 also affected cell viability and apoptosis with E2F expression in PDAC tissues correlating with expression of apoptosis and mitosis pathway genes, suggesting that E2F factors promote cell cycle regulation and tumorigenesis in PDAC cells. In conclusion, our findings show that E2F1 and E2F8 transcription factors regulate cell proliferation, survival, and migration during pancreatic carcinogenesis.

## Introduction

Pancreatic ductal adenocarcinoma (PDAC), the most prevalent pancreatic cancer (1), has a poor prognosis and low survival rate (less than 10% 5-year survival rate). Incidences are expected to rise further in the near future with projections indicating a more than 2-fold increase in the number of cases within the next decade (2–4). Low survival rates are caused by delayed diagnosis which results in initiation of treatment at an advanced stage when tumor cells have already started to invade surrounding tissues and metastasize (5, 6). A prominent reason for delayed diagnostics is the lack of PDAC biomarkers, making it a priority to discover novel genetic and biological features of PDAC to facilitate diagnosis and development of novel treatments.

Recent studies relying on data derived from animal models have shown that embryonic pancreas and PDAC development share molecular similarities. Developmental regulators normally prevent apoptosis and ensure cell proliferation of progenitor cells, however, when mis-expressed in adult cells, may cause changes in cell plasticity and promote survival of tumor cells thereby contributing to tumor formation (7). For example, PDX1 is essential for pancreas formation (8) and maintenance of adult endocrine cell function (9), however aberrant PDX1 expression in adult exocrine cells promotes formation of pancreatic neoplastic lesions and epithelial to mesenchymal transition (10). Another example is the developmental transcription factor HNF1A which is highly enriched in pancreatic cancer stem cells and HNF1A overexpression promotes tumor formation and invasiveness (11). These examples illustrate that exocrine and ductal cells undergo dedifferentiation processes similar to development when becoming carcinogenic.

Human and mouse pancreas development differ significantly, restricting the possibilities of identifying common molecular pathways between human pancreas development and tumor formation using animal models. To identify novel signaling pathways in PDAC, a comparison of RNA sequencing data from human embryonic and adult pancreas with PDAC samples was performed. In this analysis, the transcriptional regulators E2F1 and E2F8 were found to be enriched in embryonic and PDAC transcriptomes.

Previous studies have linked the E2F family of transcription factors to cell cycle regulation where E2F1 mediates both cell proliferation and TP53/p53-dependent apoptosis (12), whereas E2F1’s role in PDAC has not been studied thoroughly, with only a link to PDAC chemo-resistance being reported (13). Moreover, the function of E2F7 and E2F8 which represent a unique repressive arm of the E2F transcriptional network and control the E2F1-p53 apoptotic axis by directly binding to and repressing E2F1 transcription (14), has not been studied in PDAC so far.

Single cell sequencing analysis of human embryonic pancreas and PDAC showed that E2F1 and E2F8 were expressed in proliferating pancreatic progenitor cells and ductal cancer cells and co-expression analysis revealed that E2F expression was highly correlated with cell cycle regulation in both tissues. Functional analysis in cancer cells demonstrated that loss of E2F factors resulted in reduced colony formation, tumor cell spreading and migration, proliferation, and oncogene expression, supporting a role of these transcriptional regulators in contributing to carcinogenesis.

## Materials and Methods

### Gene expression profiling in human fetal tissues

Embryonic pancreas was obtained from terminated foetuses (7–14 gestational weeks, n = 16). Informed consent was obtained from the participating women. Ethical permission has been obtained from the regional ethics committee in Lund (Dnr2012/593, Dnr 2015/241, Dnr 2018-579).

### Bulk RNAseq

RNA was extracted and libraries generated as previously described (15). For comparison with adult tissues, the RNA-Seq expression quantification pipeline from GTEX V8 (https://github.com/broadinstitute/gtex-pipeline/) was used (16, 17). Paired-end 101 bp-long reads were aligned to the Reference Human Genome Build 38 using STAR v2.5.3a (18), annotation Gencode v26. After alignment and post-processing, expression quantification was performed using RSEM and RNASeqQC (19, 20). Normalization of library sizes was performed in edgeR by dividing counts by the library counts sum and multiplying the results by a million to obtain counts per million (CPM) values.

Fetal-specific expression was defined as ≥ 1CPM in >75% of the samples. If a gene showed expression in fetal but ≤ 1CPM in adult tissues, the gene was considered to have fetal specific expression. Comparison with expression in adult pancreas: edge-R was used to perform differential expression analysis with age and sex as covariates. Batch correction was performed by considering platform and batches using COMBAT.

Single-cell RNAseq of embryonic pancreas has been performed as previously described (15). In total, 3199 cells were obtained and included in the analysis. Unsupervised clustering was performed and visualized on the Loupe browser and validated using the Seurat R package as well as custom R scripts. Normalized counts were obtained, and correlation of E2Fs genes with all the genes expressed in at least 3 cells was performed using spearman correlation implemented in R. Significant correlated genes with a false discovery rate (FDR) lower than 5% were then studied using Reactome analysis tool to identify relevant pathways (21).

### Study population

This study is comprised of a cohort of 154 patients with pancreatic adenocarcinoma operated on without neoadjuvant therapy between 2000-2013 at the Department of Surgery, Helsinki University Hospital. Clinical data were obtained from patient records and survival data were provided by the Finnish Population Registration Centre and Statistics Finland. The median age of the patients at diagnosis was 64.8 (interquartile range 59.1-71.0) and the median length of overall survival was 2.0 years (range 0.0-13.1). The Surgical Ethics Committee of HUH approved the study protocol (Dnr. HUH 226/E6/06, extension TKM02 §66 17.4.2013). Tissue archive samples examination was done with permission from the Finnish Medicines Agency (Dnr. FIMEA/2021/006901 28.12.2021). Patients provided their written informed consent upon inclusion in the study. Patient information, samples and data were handled and stored in accordance with the Declaration of Helsinki and other local regulations.

### Preparation of tumor tissue microarrays

Paraffin blocks of tumor samples from surgical specimens fixed in formalin were collected from the archives of the Department of Pathology, University of Helsinki. Hematoxylin- and eosin-stained sections were evaluated by an experienced pathologist confirming the diagnosis. Two 1.0-mm-diameter punches were taken from each tumor block with a semiautomatic tissue microarray instrument (TMA) (Beecher Instruments, Silver Spring, MD). One section was cut from each TMA block giving two spots from each tumor sample.

### Immunohistochemistry

TMA blocks were freshly cut into 4-µm sections, fixed on slides, and dried at 37°C for 12 to 24 hours. TMA-slides were treated in a PreTreatment module (Agilent Dako) with retrieval solution pH9 (Envision Flex target retrieval solution, DM828, Agilent Dako) for 15 min at 98°C. Sections were stained with Autostainer 480S (LabVision Corp.) using Dako REAL EnVision Detection System, Peroxidase/DAB+, Rabbit/Mouse. First slides were treated with Envision Flex peroxidase-blocking reagent SM801 for 5min to block endogenous peroxidases. Then slides were incubated with E2F1 (VD301790, Thermo Fischer, 1:100 diluted in Dako REAL Antibody Diluent S2022) or E2F8 antibody (ab185727, Abcam, 1:500 diluted in Dako REAL Antibody Diluent S2022) overnight or for 60min, respectively. Subsequently all slides underwent 30-min incubation with peroxidase-conjugated EnVision Flex/HRP (SM802) rabbit/mouse (ENV) reagent. Slides were visualized using DAB chromogen (EnVision Flex DAB, DM827) for 20 min. Mayers hematoxylin (S3309, Dako) was used for counterstaining. Brain and thyroid tissue served as a positive control.

### Scoring of E2F staining in tumors

Immunoreactivity in tumor cells was interpreted independently by two researchers (T.K. and J.H), while the researchers were blinded to the clinical outcome of the patients. Negative staining was scored as 0, weakly positive as 1, moderately positive as 2, and strongly positive as 3 separately for tumor cells for both E2F1 and E2F8 (Figure S1). The highest score of each sample was considered representative for the expression in both stainings. In case of discrepancy between the observers, a consensus score was used. For statistical analysis, samples were grouped into low (negative - low), moderate and high.

Survival analysis was done by the Kaplan-Meier method and compared by the log-rank test. Overall disease-specific survival (DSS) was calculated from the day of surgery until death from PDAC or until the end of the follow-up period. A p-value of ≤ 0.05 was considered significant, and tests were two-sided. Statistical analyses were done with SPSS version 27.0 (IBM SPSS Statistics version 27.0 for Mac).

### Gene expression studies in PDAC samples

Sequencing data consisting of PDAC samples (n=24) and control samples (n=11) from Peng et al (22) were used (accession number: CRA001160) and processed as closely as possible to the parameters described previously (22). 10 cell types were identified and categorized among the PDAC samples separately according to the criteria used by Peng et al (22). These cells where re-clustered in Loupe browser with default settings to get a tSNE plot, which resulted in a total of 57383 re-clustered cells (PDAC= 41862 and control=15521). E2F expression was then filtered for, and plots were generated using the built-in function in Loupe browser. Expression correlation of E2F1 and E2F8 genes was performed as described in the “Gene expression profiling in human fetal tissues” paragraph.

### Target gene analyses

Potential E2F1 or E2F8 target genes were identified using ChiP-Atlas (http://chip-atlas.org, (23)) with binding sites located around transcription start sites (± 1 kb) being considered. Potential target genes were classified according to their average binding scores of MACS2 and compared with the list of genes significantly co-expressed with E2F1 or E2F8 in PDAC tissues (see “Gene expression studies in PDAC samples”). Overall survival and disease-free survival maps of the identified target genes were performed by using the Gene Expression Profiling Interactive Analysis (GEPIA) database (http://gepia2.cancer-pku.cn/index.html, (24)).

### Cells

Human pancreatic cancer cell lines PANC-1 (25) were obtained from the European Collection of Authenticated Cell Cultures (ECACC, 87092802) and were maintained in DMEM high glucose medium (Sigma, D6429) supplemented with 10% FBS and 1% penicillin–streptomycin. Cells were incubated at 37°C in a humidified incubator with 5% CO2 and frequently tested for mycoplasma contamination using MycoAlert™ detection kit (Lonza, LT07-118).

Cells were transfected with negative control siRNA (Invitrogen,4390843), E2F1 siRNA (Invitrogen, s4406) or E2F8 siRNA (Invitrogen, s36210) at 10 nM with RNAiMax Lipofectamine (Invitrogen, 13778150) according to the manufacturer’s guidelines and cultured for 48 hours or 7 days.

### qPCR

Total RNA was extracted using RNeasy Qiagen kit and cDNA was generated. qPCR assays were performed using StepOnePlus Real-Time PCR System. Relative gene abundance was calculated using the ΔΔCt method with GAPDH, S18, and TBP as housekeeping genes and expressed as FC to control (Table S1).

### Colony assay

To detect colony formation, cells were transfected in 24-well plates at a density of 600 cells/well. Medium was changed every 2 days. After 7 days of incubation, cells were fixed with paraformaldehyde (PFA) 4% for 15min and stained with a 0,1% crystal solution in 20% EtOH for 30min. Once dried, colonies were photographed and pictures were analyzed by ColonyArea plugin (26) with ImageJ software. Colony formation was quantified by determining the percentage of area covered by cell colonies and the cell density according to the intensity of the staining.

### Proliferation and apoptosis

Proliferation was quantified using Click-iT™ Plus EdU Cell Proliferation Kit for Imaging, Alexa Fluor™ 647 dye (Thermofisher, C10640). Cells were transfected with siRNAs in 8-well chamber slides at a density of 1×10^4^ cells/well (48h) or 3,75×10^3^ cells/well (7 days). After 48h or 7 days of incubation, cells were exposed to 10 μM of 5-ethynyl-2′-deoxyuridine (EdU) for 4h at 37°C and then fixed with PFA 4%. EdU labelling was done according to manufactureŕs instructions (Life Technologies, C10640) and nuclei were stained with Hoechst 33342. Images were acquired using a confocal microscope and quantification was done using the Cell Counter Plugin in ImageJ to manually tag and count stained cells for each colour channel and apoptotic bodies.

### Triplex assay

PANC-1 cells were transfected with siRNAs in 96-well plate at a density of 2000 cells/well (48h) or 500 cells/well (7 days). Cell viability, cytotoxicity and apoptosis were measured 48h and 7 days after the knock-down using ApoTox-Glo triplex assay kit (Promega,G6320) according to manufacturer’s instructions.

### Migration

Cell culture inserts in 24-well plate (8µm pore size; Falcon, 353097) were used for the migration assay. 7 days after transfection, cells were trypsinized and counted. 10% FBS medium was used as chemoattractant in the bottom chamber and 4×10^4^ cells diluted in 0,5% FBS medium were added in the top chamber. After 7h of incubation at 37°C, cells were fixed and stained with eosin. Cells that didńt migrate were removed from the insert using a cotton swab. Insert membrane was cut and mounted and migrated cells were imaged and counted.

### Spreading

7 days after transfection cells were trypsinized, and 4×10^4^ cells diluted in 0,5% FBS medium were cultured on coverslips coated with fibronectin (Sigma, F1141) in a 12-well plate as described by Pijuan (27). 1 hour later, cells were fixed with PFA 4% and stained with a 0,1% crystal solution. Round cells were considered unspread, while cells with cytoplasm surrounding the circumference of the nucleus were considered spread cells. Ratio was calculated as the number of spread cells divided by the total number of cells on the coverslips.

### Statistics

Statistical analyses and graphs were performed using R software with ggplot2 and ggpubr packages. Each experiment has been performed at least two times independently, with at least 3 replicates each time. Multiple comparisons between control and different siRNAs were analysed using ANOVA with Tukey’s post-hoc test. Normality and homoscedasticity of the variances were verified by checking graphically (i.e. Q-Q plot and residuals versus fits plot). Data were visualized using box plots, the central mark indicates the median and edges indicate 25th and 75th percentiles. Whiskers extend to the largest or smallest point comprised within 1.5× of the interquartile range from both edges.

## Results

### E2F1 and E2F8 are expressed in the developing human pancreas and PDAC

Expression of developmental regulators is re-initiated during tumor formation (7) and altered expression is associated with malignancy. To identify key genes in PDAC development, we compared expression of genes upregulated in PDAC (28, 29) with expression data from human embryonic pancreas. E2F1, E2F3 and E2F8 genes were expressed both in PDAC and in human embryonic pancreas anlagen (Figure 1A) and expression of E2F1, E2F2, E2F5, and E2F8 was significantly higher in embryonic than in adult pancreas (Figures 1B, Table S2). Thus, E2F1 and E2F8 expression are up-regulated in PDAC and human embryonic pancreas compared to adult tissues.

**Figure 1.**
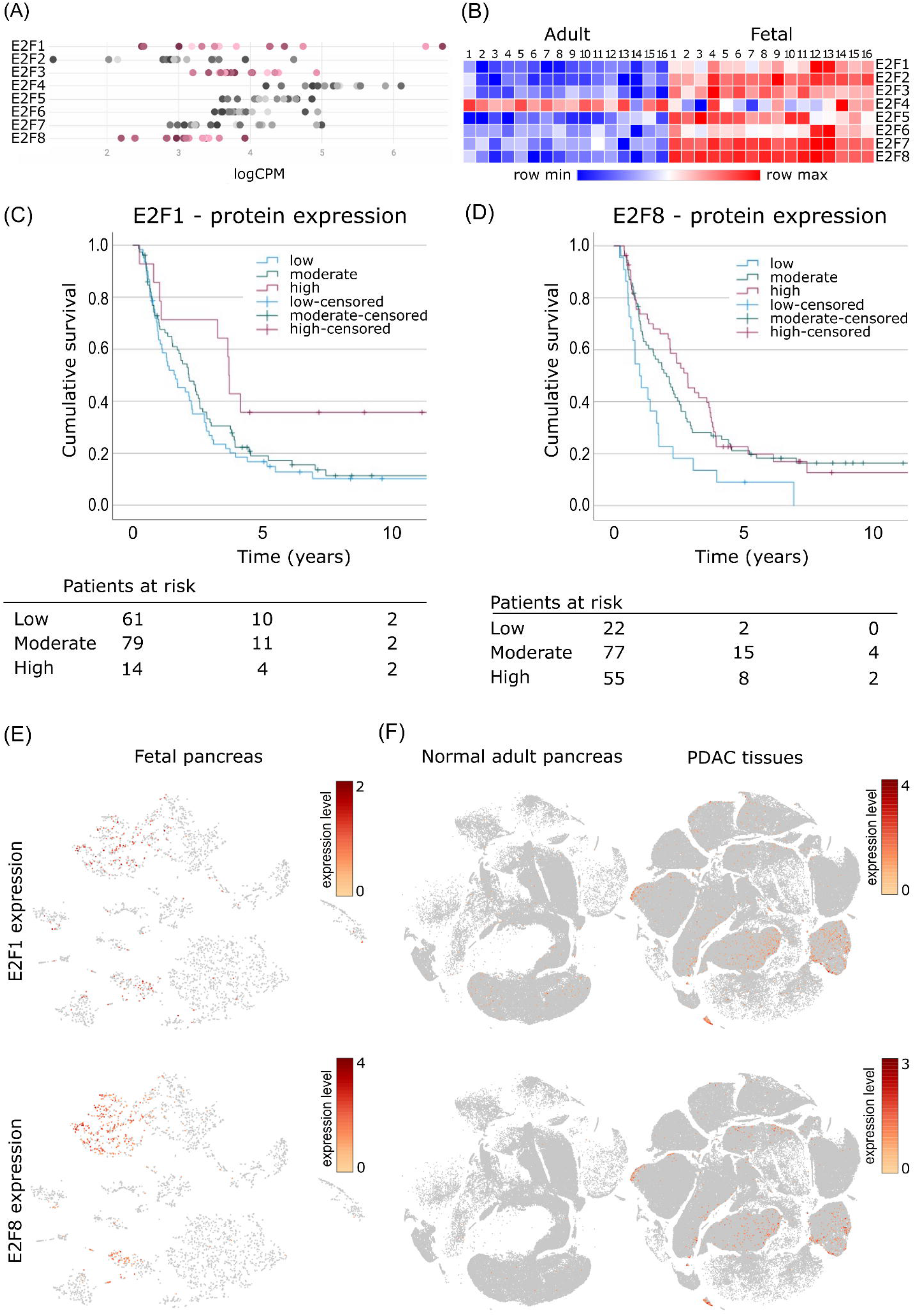
E2F gene expression in pancreas from fetal, adult, and cancer tissues. (A) Plot of E2F gene expression in fetal pancreas (7-14 weeks post-conception (PC)). X-axis shows expression in LogCPM. Pink marks E2F genes that are up regulated in PDAC both in Maós and Liús studies (27, 28). (B) Heatmap showing E2F expression in adult and fetal pancreas, with significant expression level changes for E2F1/2/4/5/8 (p-values from the differential expression analysis in Table S2). Kaplan– Meier disease-specific survival curves for PDACs according to E2F1 (C) or E2F8 (D) protein expression. (E, F) Associated expression patterns of E2F1 and E2F8 in developing pancreas from an 8-week PC embryo, in normal adult pancreas and PDAC tissues. Expression of E2F1 and E2F8 on the t-SNE embedding from Figure S1 (E) and Figure S2 (F). Any cell with E2F1 or E2F8 detected was colored according to its expression level.

TMA analysis detected E2F1 and E2F8 protein expression in 83,8 and 90,3% of analyzed PDAC samples, respectively. Interestingly expression was detected mainly in the cytoplasm (E2F1) and both cytoplasm and nucleus (E2F8) of tumor cells. Kaplan–Meier analysis revealed that E2F8 expression associates significantly with DSS, with the poorer outcome for patients with low E2F8 expression compared to moderate (p=0.015) or high E2F8 expression (p=0.004). No difference was seen between patients with moderate or high expression. 5-year DSS for patients with a low E2F8 expression was 9.1% (95% CI 0-21.1%) compared to 21.2% (95% CI 11.8-30.6%) among those with a moderate or 19.9% (95%CI 8.7-31.1%) with a high expression. Low E2F1 expression associates with worse prognosis compared high expression (p=0.028), but no difference was seen compared to moderate expression (p=0.424). No difference with statistical significance was seen between moderate and high expression (p=0.066). 5-year DSS for patients with a low E2F1 expression was 16.7% (95% CI 7.3-26.1%) compared to 19.0% (95% CI 9.0-28.0%) among those with a moderate or 35.7% (95% CI 10.6-60.8%) with a high expression. (Figure 1 C, D). No significant association with patho-clinical parameters like age, sex, T-or N-status, stage, grade, lymph node ratio or perivascular/perineural invasion were detected (Supplementary Table S3). These data suggest that variations in E2F expression influence tumor malignancy.

To assess which type of embryonic and PDAC cells expressed E2F1 and E2F8, single cell RNA sequencing analysis was performed. In human embryonic pancreas E2F1 and E2F8 expression was predominantly detected in highly proliferating pancreatic progenitor cell populations (Figure 1E, Figure S2). In PDAC, E2F1 and E2F8 expression was detected in immune and type 2 ductal cells (Figure 1F, Figure S3A, B). Type 2 ductal cells are considered highly proliferative and malignant in contrast to type 1 cells (22) and in this cell population, only 9% of the E2F1^+^ or E2F8^+^ cells co-expressed E2F1 and E2F8 (Figure S3C). These findings show that E2F1 and E2F8 mRNA expression is present during normal pancreas development and re-initiated in malignant ductal cells with aberrant expression impacting patient survival suggesting an involvement in PDAC tumor formation and maintenance.

### Loss of E2F1 and E2F8 gene expression reduces clonogenic capacity and cell proliferation

To determine if E2F genes participate in tumor biology, E2F1 and E2F8 expression was knocked down using siRNAs in the pancreatic cancer cell line PANC-1 (i.e., siE2F1 and siE2F8). This human pancreatic cancer cell line has been isolated from a pancreatic carcinoma of ductal cell origin and highly expresses E2F candidate genes (22). Knock down (KD) efficiency of E2F1 and E2F8 siRNAs was confirmed by RT-qPCR. The transfection efficiencies at 48h and 7 days after KDs were > 75 and 60% respectively (Figure S4).

To assess if siE2F1 and siE2F8 affect the capacity of tumor cells to grow we first examined the ability of KD cells to form colonies. 7 days after the transfection, the area and density of PANC-1 colonies were significantly decreased in siE2F1 and siE2F8 cells compared to the control. In mean, colony formation was reduced over 95% in siE2F1 and 70% in siE2F8 compared to control (Area and density p-values < 1e-8 for both KDs Figure 2A, B).

**Figure 2.**
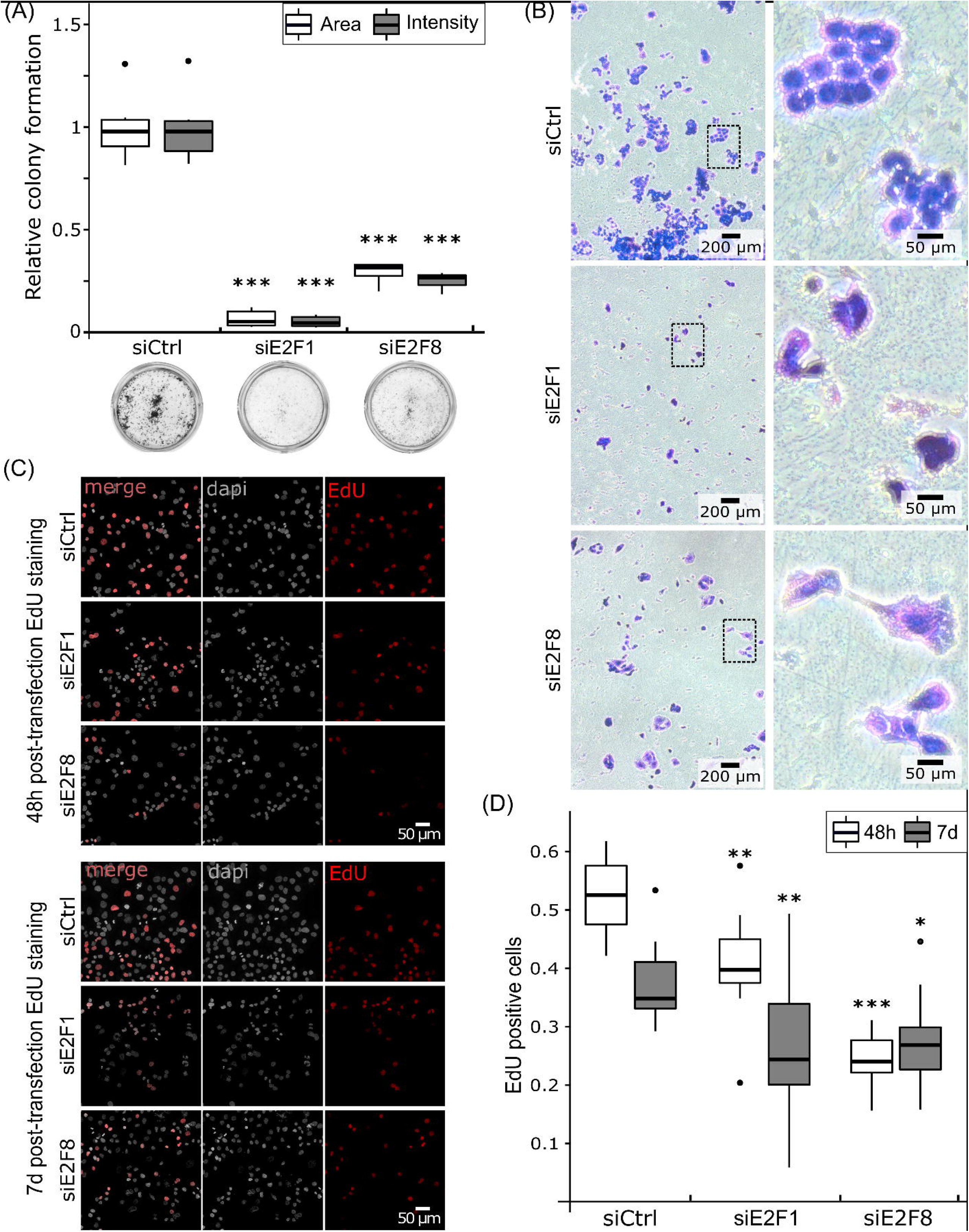
Colony formation and proliferation capacities of E2F transfected cells. Relative colony formation quantification (A) and morphological observation (B). (A) Results are shown as colony area or staining intensity percent 7 days after the knock-down compared to the negative control (transfected with scrambled RNA). (B) Colonies stained with crystal violet. (C) Dapi and EdU staining 48h or 7 days after transfection. Cells were exposed to EdU for 4h. (D) Results are shown as EdU positive cells to the total number of cells. (A, D) Measurements were derived from 2 independent experiments of three replicates per condition. Tukey’s post-hoc test significances are indicated by stars compared to the control when significant, * p < 0.05, ** p < 0.01 *** p < 0.001.

We next quantified the proliferative capacity of KD cells using an EdU incorporation assay (Figure 2C). 48h after transfection, the proportion of EdU-positive cells was significantly decreased in siE2F1 (-20%, p-value 0,0052) and siE2F8 (-54%, p-value < 1e-8) compared to the control (Figure 2D). Cell proliferation capacities were also tested 7 days post transfection to detect long term effects of siE2F1 and siE2F8. 7 days after the transfection, the proliferation capacity was still decreased in KD cells to 29% in siE2F1 (p-value 0,0055) and 26% in siE2F8 (p-value 0,019) compared to control cells. Taken together, these results suggest that E2F transcription factors regulate the cell cycle in PDAC cells.

### Loss of E2F1 expression affects cell viability and apoptosis

siE2F1 affected the clonogenic capacity to a greater extent than siE2F8 (p-value siE2F1 vs siE2F8: 0,005). To assess if this is due to an effect of E2F function on cell viability, we determined apoptosis and cell viability rate of siE2F1 and siE2F8 using Triplex assay and quantifying the apoptotic bodies. 48h after transfection, no significant differences in viability and apoptosis were observed between the KDs and control cells (Figure 3A-D). 7 days post transfection, siE2F1 significantly reduced cell viability (-60%, p-value < 1e-8), increased caspase activity (p-value 0,0001) and number of apoptotic bodies (p-value 3e-7) compared to control, while siE2F8 did not affect cell caspase activity and cell viability (Figure 3D-G). Thus, low proliferation rate and increased apoptosis contribute to the decreased clonogenic capacity of siE2F1 cells. These results suggest that reduced E2F1 but not E2F8 expression promotes apoptosis.

**Figure 3.**
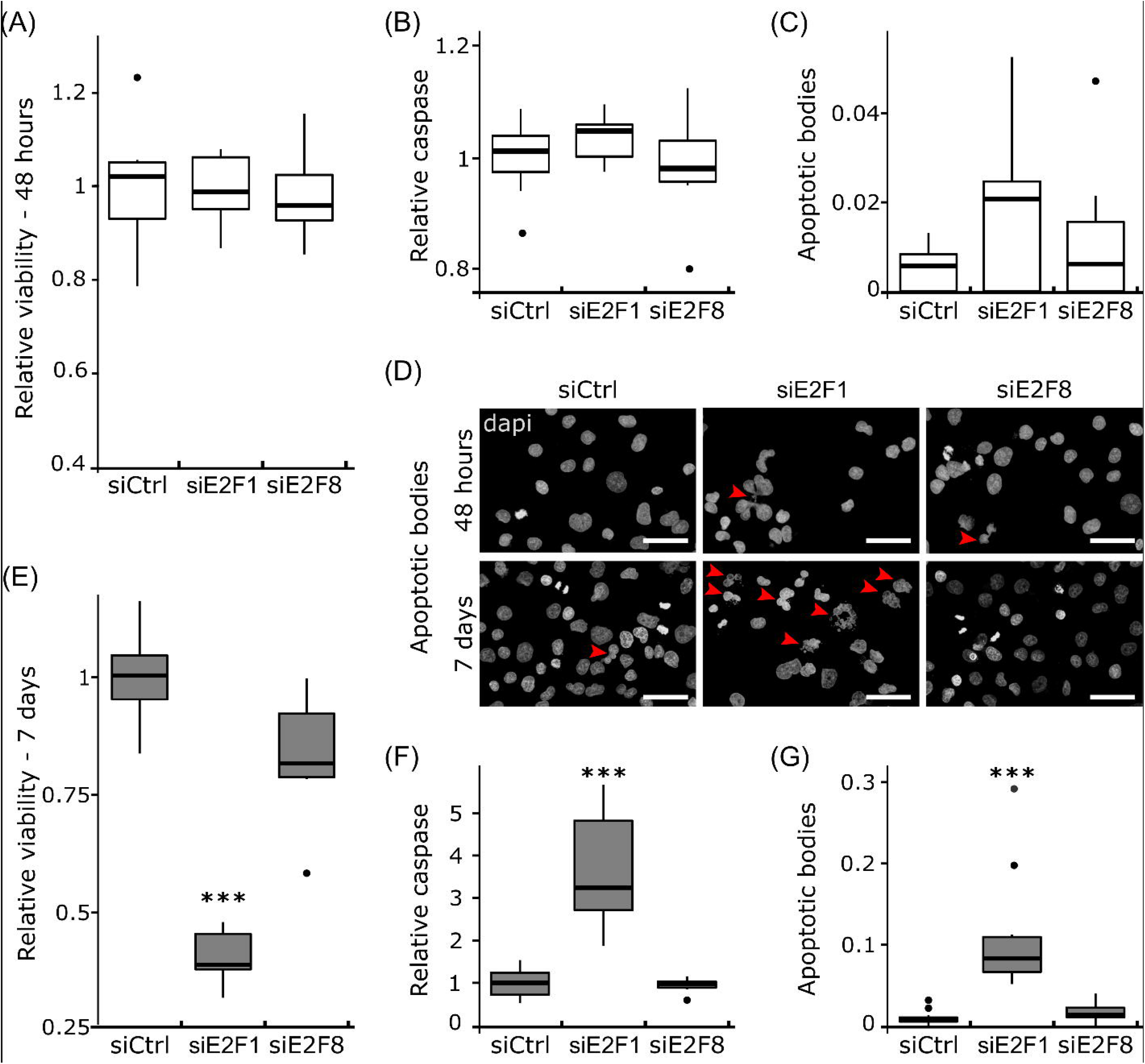
Cell viability and apoptosis 48h (A-D) and 7 days (D-G) after transfection. Cell viability 48h (A) or 7 days (E) after transfection. Results are shown as viability/cytotoxicity ratio to the negative control. Relative caspase activity 48h (B) or 7 days (F) after transfection. Results are shown as caspase-3/7 activity to the negative control. Quantification of the apoptotic bodies 48h (C) or 7 days (G) after transfection. Results are shown as counted apoptotic bodies to the total number of cells. At least 1000 cells were quantified per condition. (D) Dapi staining 48h or 7 days after transfection, red arrows show apoptotic bodies, scale bars: 50µm. (A-G) Measurements were derived from 2 independent experiments of three replicates. (A-C, E-G) Tukey’s post-hoc test significances are indicated by stars compared to the control when significant, *** p < 0.001.

### Loss of E2F1 and E2F8 impairs cell spreading and migration

Cell migration capacity is critical for tumor invasiveness. To determine if E2F1 and E2F8 regulate migration of tumor cells, cell spreading and migration assays were performed 7 days after transfection.

To study cell spreading, coverslips were coated with fibronectin, a main constituent of the tumor stroma (30) and a poor prognosis marker in PDAC (31). We observed that cell spreading was reduced to 78% and 29% in siE2F1 and siE2F8 cells, respectively (Figure 4A, B, p-values E2F1 < 1e-8, E2F8 2.67e-05) while 0.5% FBS medium was sufficient to promotes cell adhesion in control cells.

**Figure 4.**
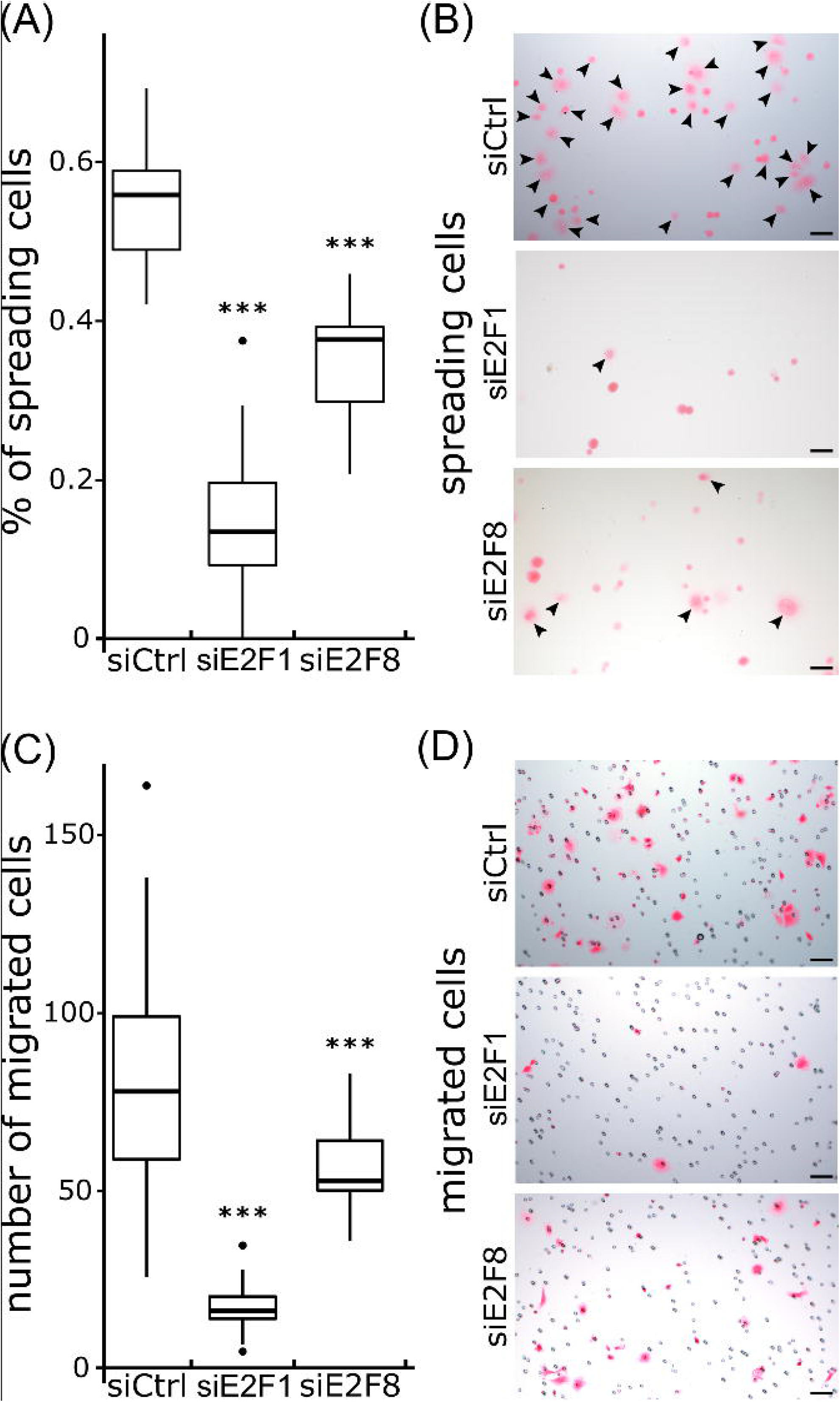
Spreading and migration capacities of PANC-1 cells 7 days after transfection. Plot showing the percentage of spreading cells (A). Cells spread on fibronectin coated plates (B). Spreading cells are indicated by a black arrow, round cells were considered to not have spread (B). Number of migrated cells per image after 7h of incubation (C) and eosin saining of the migrated cells (D). (A, C) At least 1000 cells were quantified per condition. Scale bars: 50µm. Measurements were derived from 2 independent experiments of three replicates per condition. Tukey’s post-hoc test significances are indicated by stars compared to the control when significant, *** p < 0.001.

Transwell assays were conducted to determine the migration potentials of the transfected pancreatic cancer cells. After 7 hours of incubation, control cells effectively migrated upon stimulation with chemo-attractants, while siE2F1 and siE2F8 cells had significantly reduced cell migration (Figure 4C, D, p-values siE2F1 < 1e-8, siE2F8 0,001). Thus, reduced expression of E2F1 and E2F8 reduced migration and the metastatic potential of PANC-1 cells.

### E2F transcription factors are co-expressed with genes regulating the cell cycle

The functional studies described above indicate that E2F1 and E2F8 are critical for tumor cell survival, proliferation, and migration. To identify pathways potentially regulated by E2F transcription factors and involved in carcinogenesis, unbiased co-expression analyses of E2F1 and E2F8 in single PDAC, adult and embryonic pancreatic cells were performed (Tables S4-S8).

In fetal cells, E2F8 expression significantly correlates with 2297 genes, while only one gene correlates with E2F1 expression, probably due to the lower number of E2F1^+^ cells (Tables S4-S5). In adult, 211 genes correlate with E2F1 expression (Table S6) and no gene expression correlations were obtained for E2F8 due to the low number of E2F8^+^ cells. Finally, in PDAC, unbiased co-expression analysis showed that E2F1 and E2F8 expression significantly correlates with 4989 and 612 genes, respectively (Tables S7-S8).

Only 17% of the genes significantly co-expressed with E2F1 in adult were also significantly correlated in PDAC, suggesting different target genes in normal pancreas and PDAC cells. In contrast, 51% of the genes that had significant correlation of gene expression with E2F8 in PDAC were also correlated in embryonic cells, and 56% of the genes that shared expression correlation to E2F8 in the embryo were significantly correlated to E2F1 in PDAC. These data suggest that E2F1 and E2F8 mainly act in the same gene networks during embryogenesis and PDAC. Interestingly, E2F8 correlates with the atypical repressor E2F7 in the embryo (p-value 0,0016) whereas it correlates with the activators E2F1 and E2F2 in PDAC (p-values 0,0006 and 5e-06, respectively).

In PDAC, 83% of the genes with significant positive correlation with E2F8 expression were also positively correlated with E2F1 expression, which may suggest redundant function between E2F1 and E2F8. Using Reactome analysis tools, we determined that genes co-expressed with E2F1 or E2F78 are involved in pathways regulating cell cycle and proliferation but also RNA and protein processing and translation (Table 1). These results suggest a central role of E2F1 and E2F8 in cancer cell biology.

**Table 1.** Most relevant pathways identified by Reactome using significant co-expressed genes with E2F1 or E2F8 in PDAC or embryo development. All the entities indicated are significant (p-values and FDR < 0.05)

| Most relevant pathways sorted by p-value | Nbr of entities co-expressed with |  |  | Entities in the pathway |
| --- | --- | --- | --- | --- |
|  | E2F1 in PDAC | E2F8 in PDAC | E2F8 in embryo |  |
| Formation of a pool of free 40S subunits | 100 | 53 | 75 | 106 |
| GTP hydrolysis and joining of the 60S ribosomal subunit | 110 | 54 | 79 | 120 |
| SRP-dependent cotranslational protein targeting to membrane | 109 | 54 | 79 | 119 |
| L13a-mediated translational silencing of Ceruloplasmin expression | 109 | 55 | 78 | 120 |
| Nonsense-Mediated Decay (NMD) | 110 | 53 | 80 | 124 |
| Nonsense Mediated Decay (NMD) enhanced by the Exon Junction Complex (EJC) | 110 | 53 | 80 | 124 |
| Eukaryotic Translation Initiation | 115 | 55 | 80 | 130 |
| Cap-dependent Translation Initiation | 115 | 55 | 80 | 130 |
| DNA Replication | 137 | 43 | 80 | 169 |
| Major pathway of rRNA processing in the nucleolus and cytosol | 153 | 59 | 89 | 189 |
| M Phase | 273 | 69 | 175 | 416 |
| Cell Cycle Mitotic | 392 | 129 | 256 | 596 |
| G1/S Transition | 124 | 42 | 72 | 150 |
| Cell Cycle | 485 | 148 | 314 | 734 |
| Cell Cycle Checkpoints | 214 | 57 | 140 | 279 |
| Metabolism of RNA | 567 | 130 | 309 | 829 |
| mRNA Splicing - Major Pathway | 173 | 39 | 112 | 213 |
| Processing of Capped Intron Containing Pre-mRNA | 237 | 51 | 139 | 299 |
| mRNA Splicing | 177 | 40 | 113 | 224 |
| rRNA processing in the nucleus and cytosol | 162 | 60 | 92 | 208 |
| Mitotic Metaphase and Anaphase | 177 | 43 | 106 | 250 |
| Mitotic Anaphase | 176 | 42 | 105 | 249 |
| rRNA processing | 178 | 64 | 104 | 246 |
| Translation | 261 | 71 | 124 | 339 |
| G2/M Checkpoints | 127 | 32 | 80 | 153 |
| Viral mRNA Translation | 88 | 52 | 70 | 114 |
| Nonsense Mediated Decay (NMD) independent of the Exon Junction Complex (EJC) | 94 | 53 | 71 | 101 |
| Translation initiation complex formation | 56 | 27 | 39 | 62 |
| Mitotic G1 phase and G1/S transition | 136 | 51 | 80 | 174 |
| Peptide chain elongation | 89 | 52 | 70 | 97 |
| SARS-CoV-1 modulates host translation machinery | 36 | 25 | 32 | 41 |
| Eukaryotic Translation Elongation | 92 | 53 | 72 | 102 |
| rRNA processing in the nucleus and cytosol | 162 | 60 | 92 | 208 |
| Eukaryotic Translation Termination | 92 | 52 | 70 | 106 |
| Mitotic prometaphase | 132 | 38 | 100 | 211 |

In addition, E2F1 expression specifically correlates with genes involved in the regulation of apoptosis and mitochondrial translation pathways (Table 2), suggesting a more prominent role in PDAC regulation.

**Table 2.**
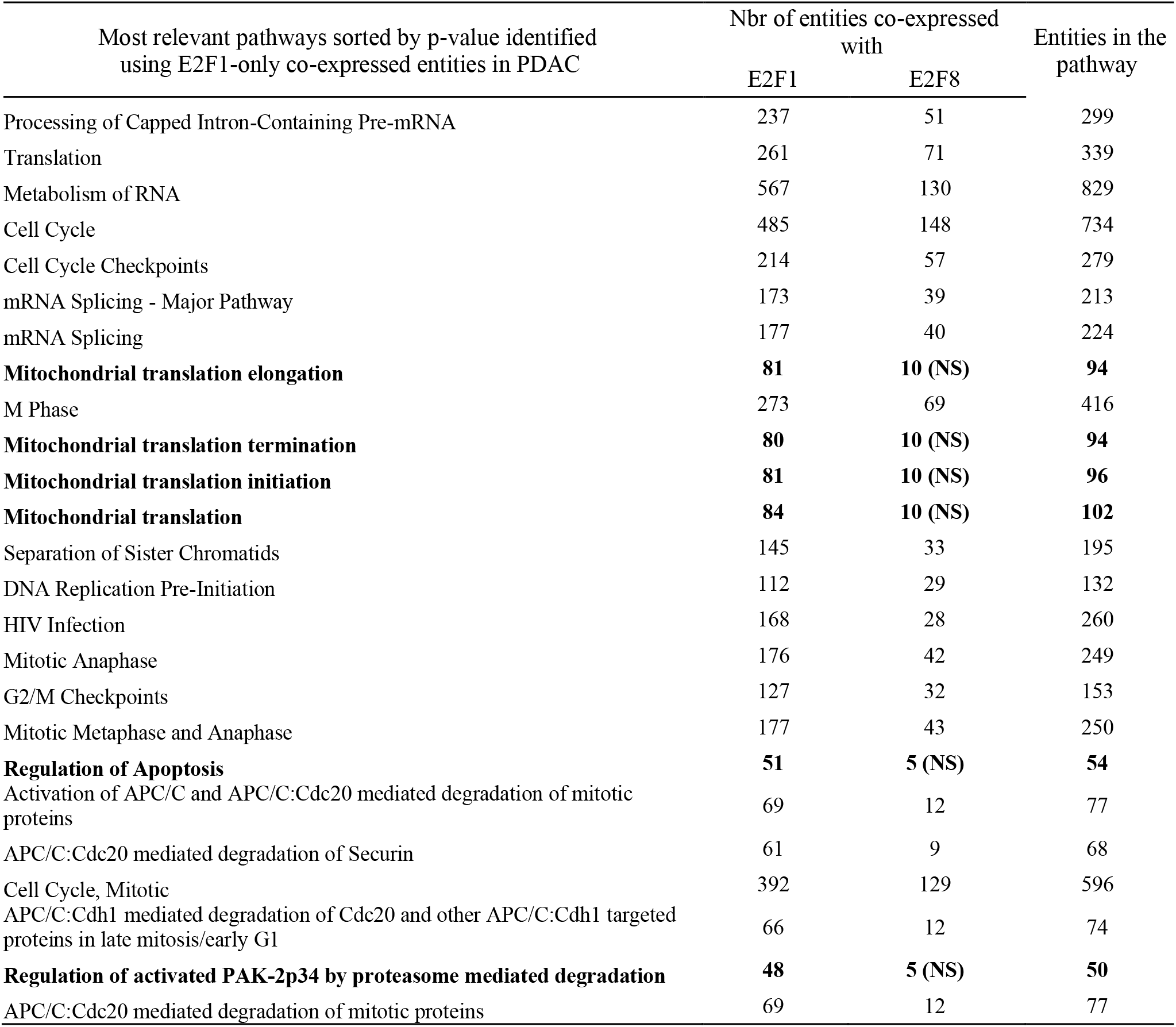
Most relevant pathways identified by Reactome using the genes significantly co-expressed only with E2F1 in PDAC. Number of entities were then identified using the full significant gene list for E2F1 or E2F8 in PDAC. In bold, the E2F1-specific pathways. All the entities are significant (p-values and FDR < 0.05) when NS isńt indicated

### E2F1 and E2F8 transcription factors can directly regulate known oncogenes

A comparison of genes co-expressed with E2F1 or E2F8 in PDAC (Tables S7-S8) with published ChiP-seq results (ChIP-Atlas) was performed to obtain a list of potential direct E2F target genes in E2F^+^ PDAC cells. Once classified by Reactome pathways (Tables S9-S10), we observed that most of these genes were involved in pathways regulating carcinogenesis like cell cycle, DNA replication, gene expression and programmed cell death.

A thorough literature review and GEPIA2 analysis showed that most of these genes were already known as potential oncogene and/or linked to poor survival in PDAC but few of them have been shown to be regulated by E2F transcription factors (Figure 5A, Tables S11-S12).

**Figure 5.**
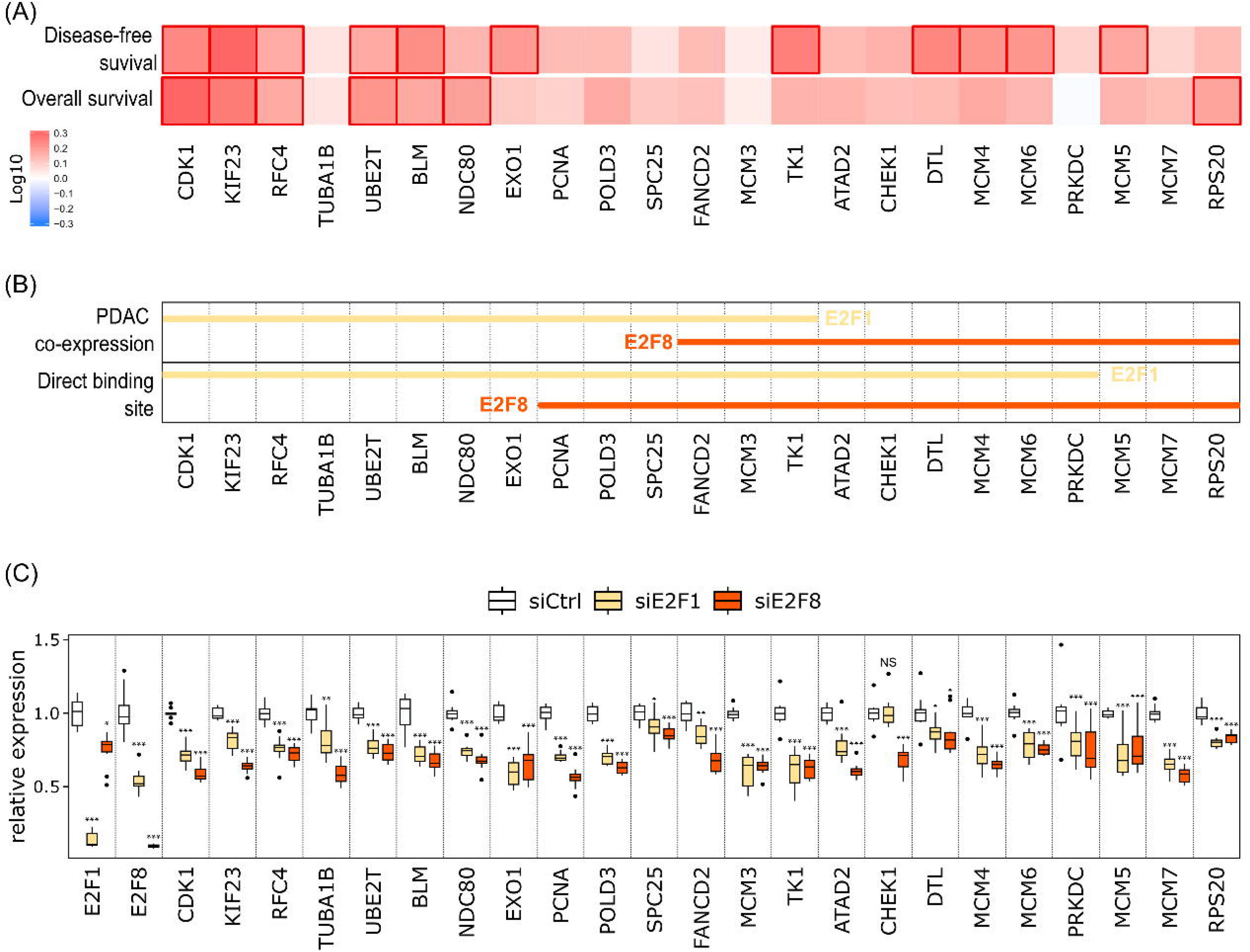
Regulation of known oncogenes by E2F1 and E2F8. (A) Disease-free survival and overall survival heatmaps according to the expression of direct E2F target genes in PDAC. The red blocks denote higher risks. The rectangles with frames mean the significant unfavorable results in prognostic analyses. (B) For each oncogene, the lines indicate that the gene is co-expressed with E2F1 (yellow) or E2F8 (orange) in PDAC tissues and contains a direct binding site for E2F1 (yellow) or E2F8 (orange). (C) qPCR analyses showing the expression of 25 oncogenes 48h after E2F1 or E2F8 KD. Significance was determined for each gene by a 1-way ANOVA and Tukey’s post-hoc test. NS p > 0.05, * p < 0.05, ** p < 0.01 *** p < 0.001.

To validate our analysis, we selected 25 oncogenes identified as potential E2F1 and/or E2F8 targets (Figure 5B) and assessed if their expression was impaired in E2F KD cells. 48h post siE2F1 or siE2F8 transfection, all the selected genes were significantly downregulated in siE2F1 and/or siE2F8 cells compared to controls (Figure 5C). These results suggest that E2F1 and E2F8 directly regulate the expression of numerous known oncogenes in PDAC tumors and thus act as master regulators of pancreatic cancer biology.

## Discussion

Great progress has been made in cancer treatment with overall survival rates increasing for many different cancer types (32). However, the diagnosis and treatment of pancreatic cancer has not been improved significantly during the past years. Immuno- and chemotherapy have limited success and only after total resection of tumor tissue. Thus, great efforts have been made to identify novel biomarkers and drugable targets. This may be facilitated by novel expression data sets from PDAC tissue. Embryonic progenitor and PDAC cells share many features like cell plasticity, survival and proliferation potential suggesting that comparison between these cell types may lead to the identification of important genes for pancreatic tumorigenesis. Here we compared single and bulk gene expression from PDAC with expression data from human embryonic pancreas to identify novel PDAC genes that are present in embryonic pancreatic progenitor and tumor cells. Using this expression correlation approach, we identified E2F genes to be highly expressed in embryonic pancreas and PDAC.

In the developing pancreas, E2F1 and E2F8 are expressed in proliferating pancreatic progenitor cells suggesting a role in cell cycle regulation in human embryos. This is consistent with previous findings showing that E2F1 and E2F8 are present in developing Pdx1+ pancreatic progenitor cells (33, 34) and that E2F1 is essential for regulating proliferation in the mouse pancreas anlage (34). E2F1 and E2F8 expression is also enriched in PDAC and the majority of E2F1 and E2F8 producing cells are malignant ductal cells suggesting that E2F transcription factors may also participate in PDAC tumor formation similar to what has been observed in tumorigenesis and treatment resistance of colon, breast and cervical cancer (35–37). Previous reports showed that high expression of mainly E2F1 is predictive of a poor clinical outcome in PDAC patients (38, 39), while our analysis of E2F gene expression in PDAC tissue showed that low E2F8 expression was associated with poor disesease specific patient survival. Interestingly, E2F8 expression is mainly detected in the immunogenic PDAC subtype (40) and correlates with all the signature genes of the classical subtype (24, 41) but not with the genes linked to the basal subtype. These observations suggest that the low E2F8 expressing group in our survival group may identify patients with basal PDAC which have a much shorter DSS rate, while patients with classical PDAC have moderate to high E2F8 expression.

E2F1 is a cell cycle regulator which has been associated indirectly with PDAC by studies describing potential upstream (42) or downstream (43) target genes that play a significant role in PDAC biology. By studying direct E2F1 down-regulation, we observed that E2F1 was not only regulating proliferation as proposed previously, but also apoptosis and cell migration. In normal tissues, a negative feedback loop regulates E2F signaling, E2F1 inducing expression of the repressor E2F8 (44). Few studies followed E2F8 function in cancer, and so far, no potential E2F8 function in PDAC has been reported. E2F8 has a dual role in cancer, as it can act as tumor suppressor (45) or proto-oncogene (46, 47) depending on cancer type. Here we showed that in pancreas tumorigenesis, E2F8 may regulate not only the cell cycle but also migration.

KD of E2F1 and E2F8 in the PDAC cell line PANC-1 resulted in reduced colony formation, with E2F1 having a significantly stronger effect, suggesting a more prominent role for E2F1 in PDAC. This is consistent with our findings that E2F1, but not E2F8 KD induced tumor cell apoptosis. This observation is in contrast to previous studies which have shown that E2F1 is essential for inducing apoptosis in response to genotoxic treatment in early tumor cells (48), however elevated E2F1 expression results in resistance to apoptosis in late stage tumor cells (reviewed in (49)) which suggests a dual role of this gene in regulating cell viability/apoptosis of PDAC cells. PANC-1 cell proliferation was impaired upon E2F1 and E2F8 KDs suggesting that E2F factors promote cell cycle regulation in PDAC cells. A finding that is further supported by expression correlation of E2F1 and E2F8 with genes involved in cell cycle regulation and mitosis in PDAC tissues.

No significant correlation between E2F1 or E2F8 with EMT marker genes was found and E2F1 and E2F8 knock down did not induce mesenchymal marker gene expression in PANC-1 cells (data not shown) suggesting that the reduced cell spreading/migration observed upon E2F1 and E2F8 knock down is most likely due to E2F regulating cell proliferation and survival.

E2F1 and E2F8 mainly positively correlate with the same genes in PDAC tissues, suggesting similar and redundant functions, as observed experimentally in pancreatic cancer cells. Our co-expression analyses showed that the same pathways were correlating with E2F1 and E2F8 expression in PDAC and embryonic tissues while it is known that in healthy tissues, E2F8 and E2F1 have antagonistic functions (14). Interestingly, in embryonic tissues, E2F8 expression correlates with the atypical repressor E2F7 whereas in PDAC E2F8 expression correlates with the transcriptional activators E2F1 and E2F2. Morgunova et al., showed that both atypical and typical E2Fs can bind to similar DNA sequences (50). Our data suggest that E2F8 may regulate the same pathways during PDAC and embryonic development but acts differently according to its co-factors.

Links between E2F genes and mitochondrial functions are now well studied and revealed a role of E2F1 as regulator of metabolism (51–53). In pancreatic cancer, it has been shown that E2F1 may induces aerobic glycolysis (53), here we link E2F1 expression in PDAC with expression of genes involved in the mitochondrial translation machinery. Interestingly, inhibition of mitochondrial functions is explored as a promising cancer therapeutic strategy in different tumor types, and some clinical trials are ongoing (54).

Using available ChiP-seq data, we identified genes containing E2F binding sites that are co-expressed with E2F1 or E2F8 in PDAC. Most of these genes are known to be over-expressed in PDAC and act in numerous oncogenic pathways. For example, ATAD2 expression is linked to cell invasion, migration and gemcitabine resistance (55) and overexpression of TUBA1B, NDC80, RCF4 and MCM proteins is associated with poorer outcome (56–59). Using E2F1 and E2F8 knock down cells, we confirmed that loss of E2F impaired expression of the identified targets genes. These results suggest that E2F1 and E2F8 over-expression in PDAC induce the expression of numerous oncogenes involved in the regulation of the cell cycle as PCNA (60) and CDK1 (61) which affect the patient prognosis.

## Conclusion

In summary our findings illustrate that E2F1 and E2F8 are expressed in pancreatic progenitor and PDAC cells, where they contribute to tumor cell expansion by regulation of cell proliferation, viability and cell migration making these genes attractive therapeutic targets and potential prognostic markers for pancreatic cancer.

## Author Contributions

Conceptualization, LBB and IA; methodology, LBB, SB, JA, OA, GH, RBP, TK, JH, CH, HS; formal analysis, LBB; investigation, LBB, IA, TK, JH, CH, HS; data curation, RBP, JA, OA, GH; writing original draft preparation, LBB; writing review and editing, IA; visualization, LBB, RBP, JA, TK.; supervision, IA; funding acquisition, LBB and IA. All authors have read and agreed to the published version of the manuscript.

## Data Availability Statement

The sequencing data related to fetal samples generated for this study can be found in the LUDC repository (www.ludc.lu.se/resources/repository) under the following accession numbers and are available upon reasonable request: LUDC2021.10.12 (bulk RNA sequencing data from fetal pancreata), and LUDC2021.10.18 (single cell RNA sequencing data from fetal pancreas).

## Abbreviations

PDAC: pancreatic ductal adenocarcinoma
CPM: counts per million
TMA: tissue microarray
FDR: false discovery rate
PC: post-conception
DSS: disease-specific survival
EdU: 5-ethynyl-2′-deoxyuridine
ChiP-seq: Chromatin Immunoprecipitation Sequencing
KD: knock down
EMT: epithelial–mesenchymal transition

## Supporting information

Legends for supplemental files

---

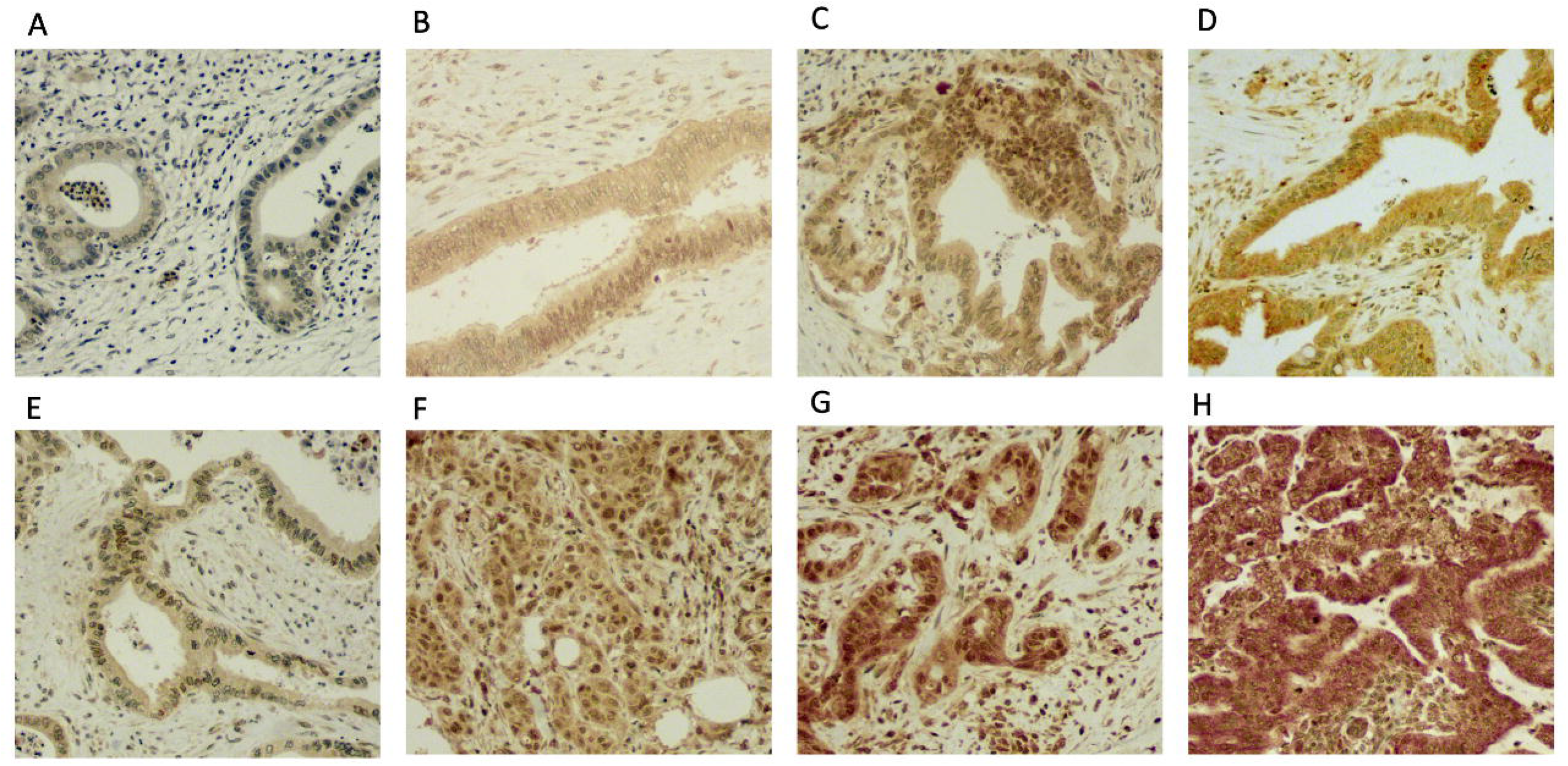

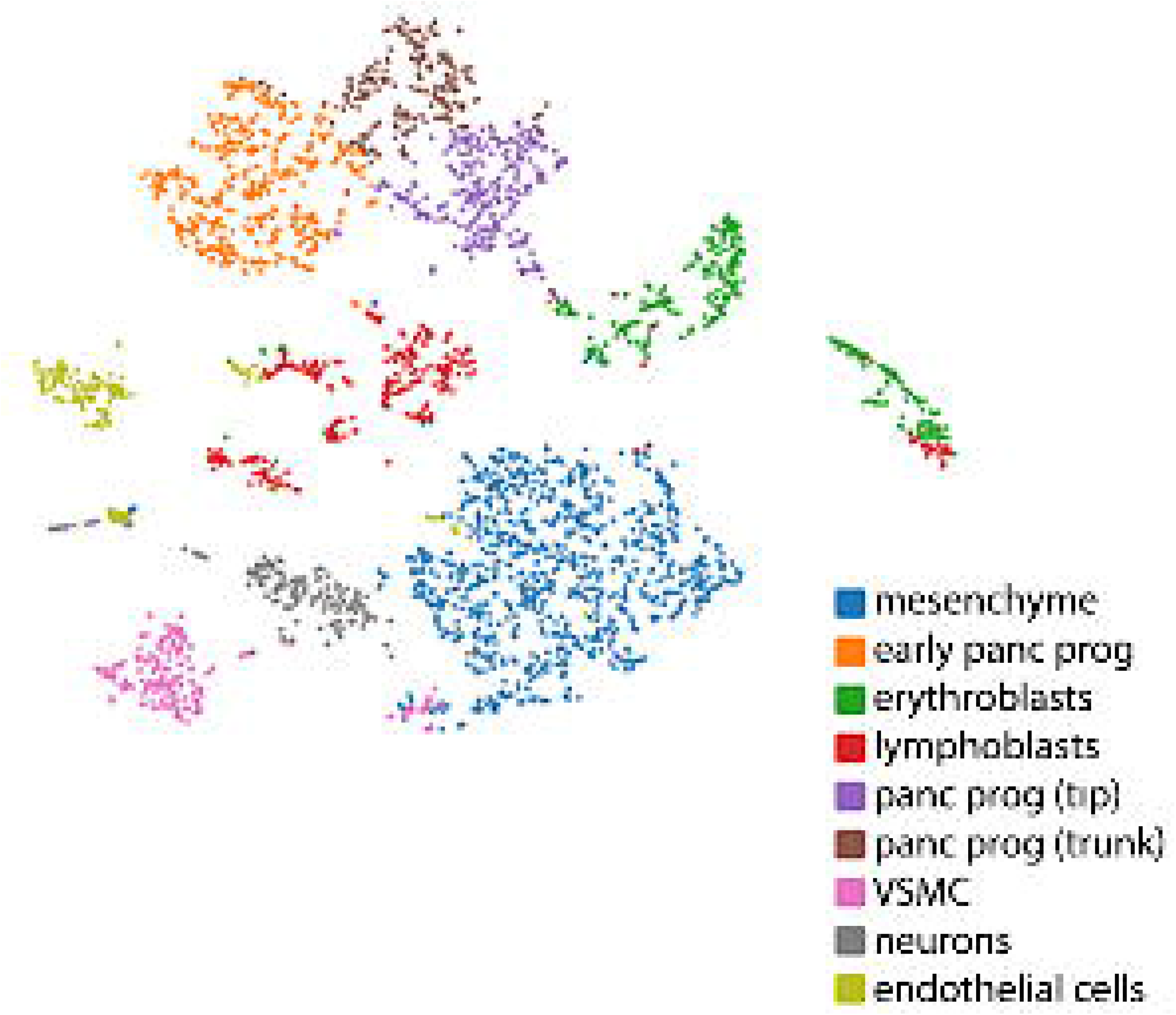

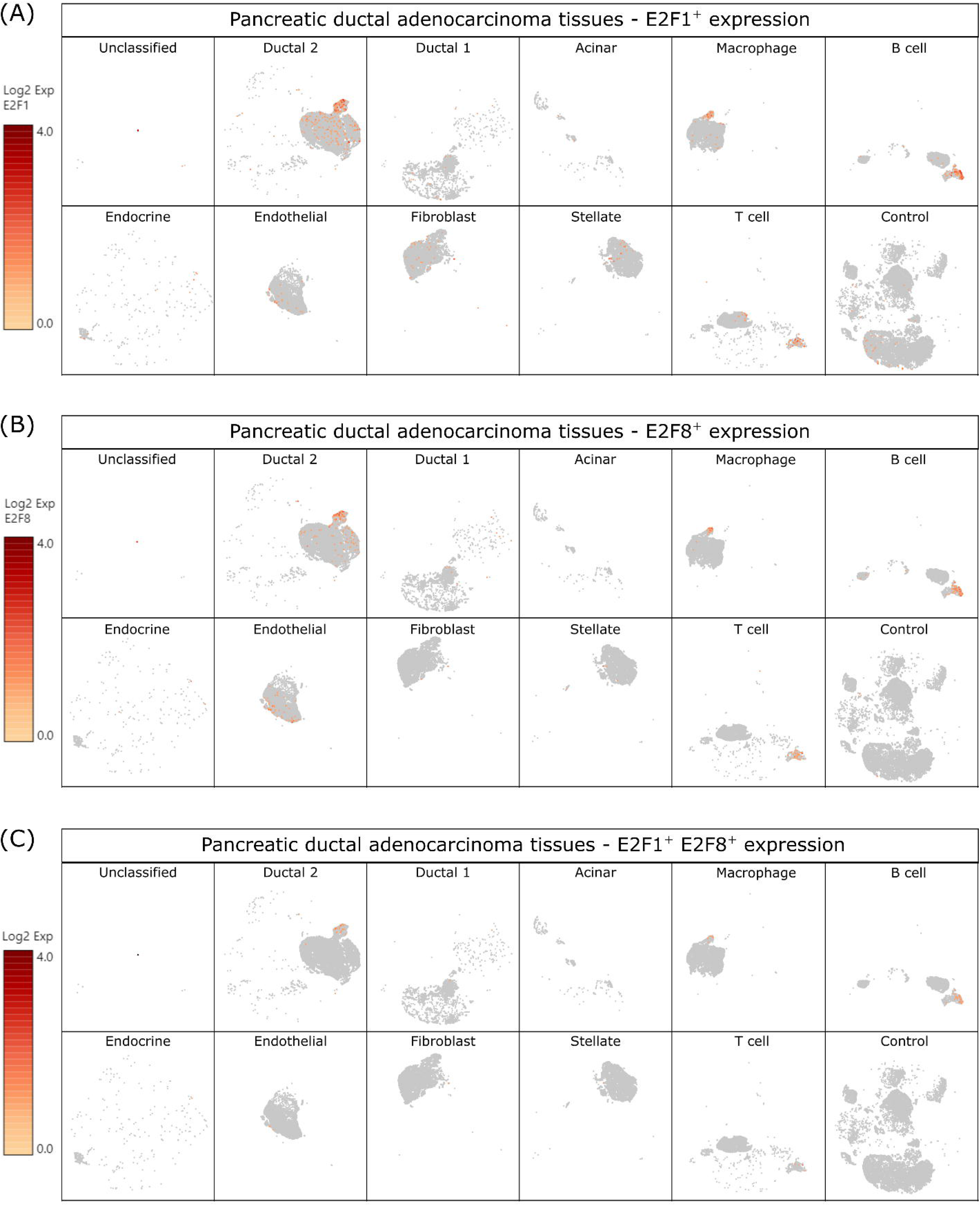

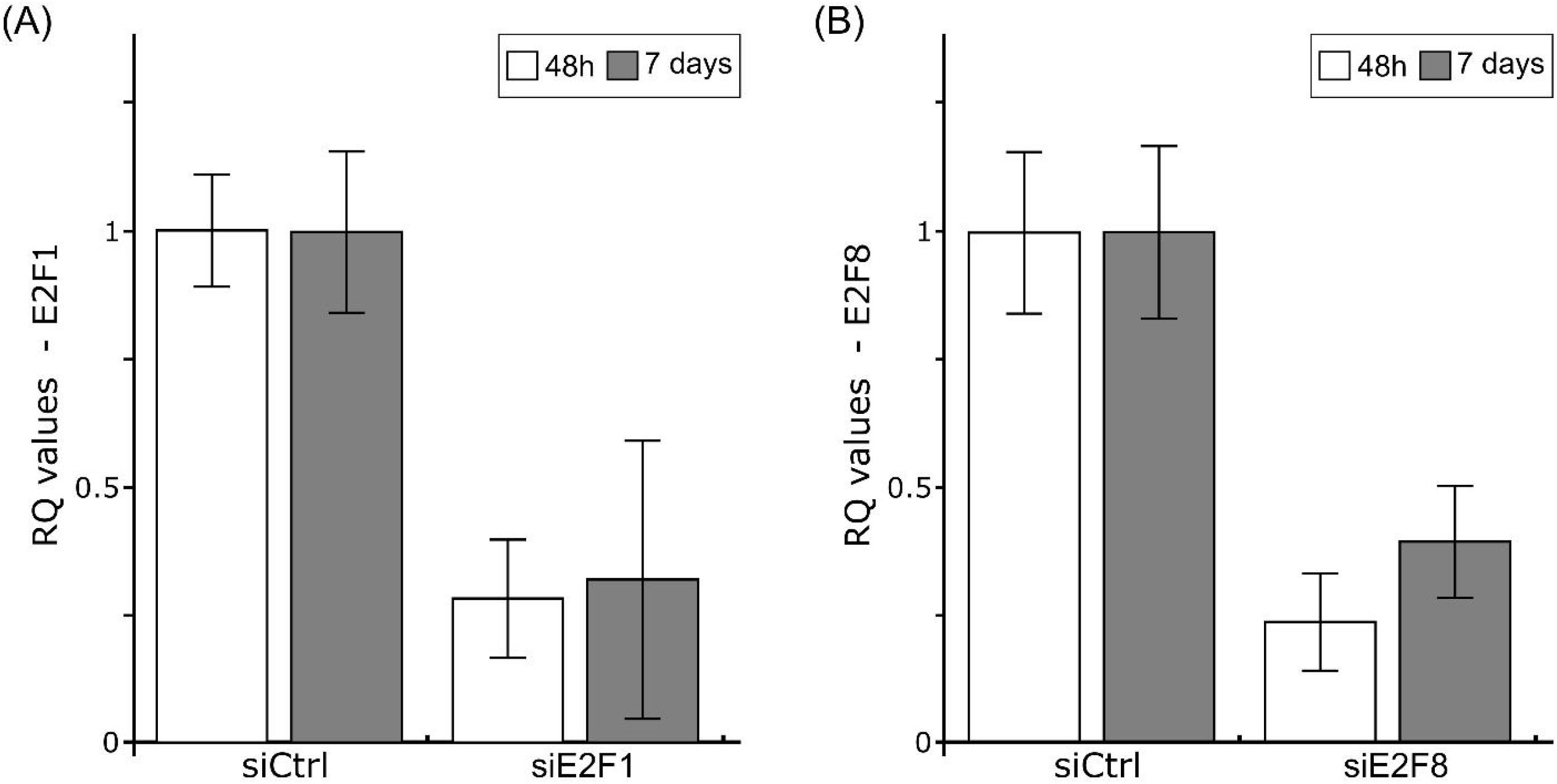

## References

1. Kleeff J, Korc M, Apte M, La Vecchia C, Johnson CD, Biankin AV, et al. Pancreatic Cancer. Nature Reviews Disease Primers (2016) 2(1):1–22. doi: 10.1038/nrdp.2016.22.

2. Hu JX, Lin YY, Zhao CF, Chen WB, Liu QC, Li QW, et al. Pancreatic Cancer: A Review of Epidemiology, Trend, and Risk Factors. World Journal of Gastroenterology (2021) 27(27):4298-. doi: 10.3748/WJG.V27.I27.4298.

3. Quante AS, Ming C, Rottmann M, Engel J, Boeck S, Heinemann V, et al. Projections of Cancer Incidence and Cancer-Related Deaths In germany by 2020 and 2030. Cancer Medicine (2016) 5(9):2649–56. doi: 10.1002/cam4.767.

4. Rahib L, Smith BD, Aizenberg R, Rosenzweig AB, Fleshman JM, Matrisian LM. Projecting Cancer Incidence and Deaths to 2030: The Unexpected Burden of Thyroid, Liver, and Pancreas Cancers in the United States. Cancer Research (2014) 74(11):2913–21. doi: 10.1158/0008-5472.CAN-14-0155.

5. Kamisawa T, Wood LD, Itoi T, Takaori K. Pancreatic Cancer. The Lancet (2016) 388(10039):73–85. doi: 10.1016/S0140-6736(16)00141-0.

6. Siegel RL, Miller KD, Fuchs HE, Jemal A. Cancer Statistics, 2021. CA: A Cancer Journal for Clinicians (2021) 71(1):7–33. doi: 10.3322/caac.21654.

7. Naqvi AAT, Hasan GM, Hassan MI. Investigating the Role of Transcription Factors of Pancreas Development in Pancreatic Cancer. Pancreatology (2018) 18(2):184–90. doi: 10.1016/j.pan.2017.12.013.

8. Kaneto H, Matsuoka TA, Miyatsuka T, Kawamori D, Katakami N, Yamasaki Y, et al. Pdx-1 Functions as a Master Factor in the Pancreas. Frontiers in Bioscience (2008) 13(16):6406–20. doi: 10.2741/3162/PDF.

9. Zhu Y, Liu Q, Zhou Z, Ikeda Y. Pdx1, Neurogenin-3, and Mafa: Critical Transcription Regulators for Beta Cell Development and Regeneration. Stem Cell Research and Therapy (2017) 8(1):1–7. doi: 10.1186/s13287-017-0694-z.

10. Roy N, Takeuchi KK, Ruggeri JM, Bailey P, Chang D, Li J, et al. Pdx1 Dynamically Regulates Pancreatic Ductal Adenocarcinoma Initiation and Maintenance. Genes and Development (2016) 30(24):2669–83. doi: 10.1101/gad.291021.116.

11. Abel EV, Goto M, Magnuson B, Abraham S, Ramanathan N, Hotaling E, et al. Hnf1a Is a Novel Oncogene That Regulates Human Pancreatic Cancer Stem Cell Properties. eLife (2018) 7. doi: 10.7554/eLife.33947.

12. Hershko T, Chaussepied M, Oren M, Ginsberg D. Novel Link between E2f and P53: Proapoptotic Cofactors of P53 Are Transcriptionally Upregulated by E2f. Cell Death Differ (2005) 12(4):377–83. doi: 10.1038/sj.cdd.4401575.

13. Ropolo A, Catrinacio C, Renna FJ, Boggio V, Orquera T, Gonzalez CD, et al. A Novel E2f1-Ep300-Vmp1 Pathway Mediates Gemcitabine-Induced Autophagy in Pancreatic Cancer Cells Carrying Oncogenic Kras. Frontiers in Endocrinology (2020) 11:411-. doi: 10.3389/fendo.2020.00411.

14. Li J, Ran C, Li E, Gordon F, Comstock G, Siddiqui H, et al. Synergistic Function of E2f7 and E2f8 Is Essential for Cell Survival and Embryonic Development. Dev Cell (2008) 14(1):62–75. doi: 10.1016/j.devcel.2007.10.017.

15. Bsharat S, Monni E, Singh T, Johansson JK, Achanta K, Bertonnier-Brouty L, et al. Mafb-Dependent Neurotransmitter Signaling Promotes Beta Cell Migration in the Developing Pancreas. Development (2023) 150(6). Epub 20230327. doi: 10.1242/dev.201009.

16. Consortium GT, Laboratory DA, Coordinating Center-Analysis Working G, Statistical Methods groups-Analysis Working G, Enhancing Gg, Fund NIHC, et al. Genetic Effects on Gene Expression across Human Tissues. Nature (2017) 550(7675):204–13. doi: 10.1038/nature24277.

17. Gamazon ER, Segre AV, van de Bunt M, Wen X, Xi HS, Hormozdiari F, et al. Using an Atlas of Gene Regulation across 44 Human Tissues to Inform Complex Disease- and Trait-Associated Variation. Nat Genet (2018) 50(7):956–67. Epub 20180628. doi: 10.1038/s41588-018-0154-4.

18. Dobin A, Davis CA, Schlesinger F, Drenkow J, Zaleski C, Jha S, et al. Star: Ultrafast Universal Rna-Seq Aligner. Bioinformatics (2013) 29(1):15–21. Epub 20121025. doi: 10.1093/bioinformatics/bts635.

19. DeLuca DS, Levin JZ, Sivachenko A, Fennell T, Nazaire MD, Williams C, et al. Rna-Seqc: Rna-Seq Metrics for Quality Control and Process Optimization. Bioinformatics (2012) 28(11):1530–2. Epub 20120425. doi: 10.1093/bioinformatics/bts196.

20. Li B, Dewey CN. Rsem: Accurate Transcript Quantification from Rna-Seq Data with or without a Reference Genome. BMC Bioinformatics (2011) 12:323. Epub 20110804. doi: 10.1186/1471-2105-12-323.

21. Gillespie M, Jassal B, Stephan R, Milacic M, Rothfels K, Senff-Ribeiro A, et al. The Reactome Pathway Knowledgebase 2022. Nucleic Acids Res (2022) 50(D1):D687–D92. doi: 10.1093/nar/gkab1028.

22. Peng J, Sun B-F, Chen C-Y, Zhou J-Y, Chen Y-S, Chen H, et al. Single-Cell Rna-Seq Highlights Intra-Tumoral Heterogeneity and Malignant Progression in Pancreatic Ductal Adenocarcinoma. Cell Research (2019) 29(9):725–38. doi: 10.1038/s41422-019-0195-y.

23. Oki S, Ohta T, Shioi G, Hatanaka H, Ogasawara O, Okuda Y, et al. Chip-Atlas: A Data-Mining Suite Powered by Full Integration of Public Chip-Seq Data. EMBO Rep (2018) 19(12). Epub 20181109. doi: 10.15252/embr.201846255.

24. Tang Z, Kang B, Li C, Chen T, Zhang Z. Gepia2: An Enhanced Web Server for Large-Scale Expression Profiling and Interactive Analysis. Nucleic Acids Res (2019) 47(W1):W556–W60. doi: 10.1093/nar/gkz430.

25. Lieber M, Mazzetta J, Nelson-Rees W, Kaplan M, Todaro G. Establishment of a Continuous Tumor-Cell Line (Panc-1) from a Human Carcinoma of the Exocrine Pancreas. Int J Cancer (1975) 15(5):741–7. doi: 10.1002/ijc.2910150505.

26. Guzmán C, Bagga M, Kaur A, Westermarck J, Abankwa D. Colonyarea: An Imagej Plugin to Automatically Quantify Colony Formation in Clonogenic Assays. PloS one (2014) 9(3). doi: 10.1371/journal.pone.0092444.

27. Pijuan J, Barceló C, Moreno DF, Maiques O, Sisó P, Marti RM, et al. In Vitro Cell Migration, Invasion, and Adhesion Assays: From Cell Imaging to Data Analysis. Frontiers in Cell and Developmental Biology (2019) 7(JUN):107-. doi: 10.3389/FCELL.2019.00107/BIBTEX.

28. Mao Y, Shen J, Lu Y, Lin K, Wang H, Li Y, et al. Rna Sequencing Analyses Reveal Novel Differentially Expressed Genes and Pathways in Pancreatic Cancer. Oncotarget (2017) 8(26):42537–47. doi: 10.18632/oncotarget.16451.

29. Liu XS, Gao Y, Liu C, Chen XQ, Zhou LM, Yang JW, et al. Comprehensive Analysis of Prognostic and Immune Infiltrates for E2f Transcription Factors in Human Pancreatic Adenocarcinoma. Frontiers in Oncology (2021) 10:3297-. doi: 10.3389/fonc.2020.606735.

30. Hu D, Ansari D, Zhou Q, Sasor A, Said Hilmersson K, Andersson R. Stromal Fibronectin Expression in Patients with Resected Pancreatic Ductal Adenocarcinoma. World J Surg Oncol (2019) 17(1):29. Epub 20190208. doi: 10.1186/s12957-019-1574-z.

31. Amrutkar M, Aasrum M, Verbeke CS, Gladhaug IP. Secretion of Fibronectin by Human Pancreatic Stellate Cells Promotes Chemoresistance to Gemcitabine in Pancreatic Cancer Cells. BMC Cancer (2019) 19(1):596. Epub 20190617. doi: 10.1186/s12885-019-5803-1.

32. Siegel RL, Miller KD, Fuchs HE, Jemal A. Cancer Statistics, 2022. CA: A Cancer Journal for Clinicians (2022) 72(1):7–33. doi: 10.3322/CAAC.21708.

33. Yu XX, Qiu WL, Yang L, Zhang Y, He MY, Li LC, et al. Defining Multistep Cell Fate Decision Pathways During Pancreatic Development at Single-Cell Resolution. The EMBO Journal (2019) 38(8):e100164. doi: 10.15252/embj.2018100164.

34. Kim SY, Rane SG. The Cdk4-E2f1 Pathway Regulates Early Pancreas Development by Targeting Pdx1+ Progenitors and Ngn3+ Endocrine Precursors. Development (2011) 138(10):1903–12. doi: 10.1242/dev.061481.

35. Xu ZH, Qu H, Ren YY, Gong ZZ, Ri HJ, Chen X. An Update on the Potential Roles of E2f Family Members in Colorectal Cancer. Cancer Management and Research (2021) 13:5509-. doi: 10.2147/CMAR.S320193.

36. Ye L, Guo L, He Z, Wang X, Lin C, Zhang X, et al. Upregulation of E2f8 Promotes Cell Proliferation and Tumorigenicity in Breast Cancer by Modulating G1/S Phase Transition. Oncotarget (2016) 7(17):23757-. doi: 10.18632/ONCOTARGET.8121.

37. Eoh KJ, Kim HJ, Lee JW, Kim LK, Park SA, Kim HS, et al. E2f8 Induces Cell Proliferation and Invasion through the Epithelial–Mesenchymal Transition and Notch Signaling Pathways in Ovarian Cancer. International Journal of Molecular Sciences (2020) 21(16):1–16. doi: 10.3390/IJMS21165813.

38. Luo L, Zhang G, Wu T, Wu G. Prognostic Value of E2f Transcription Factor Expression in Pancreatic Adenocarcinoma. Medical Science Monitor: International Medical Journal of Experimental and Clinical Research (2021) 27:e933443–1. doi: 10.12659/MSM.933443.

39. Lan W, Bian B, Xia Y, Dou S, Gayet O, Bigonnet M, et al. E2f Signature Is Predictive for the Pancreatic Adenocarcinoma Clinical Outcome and Sensitivity to E2f Inhibitors, but Not for the Response to Cytotoxic-Based Treatments. Scientific Reports (2018) 8(1):8330-. doi: 10.1038/S41598-018-26613-Z.

40. Bailey P, Chang DK, Nones K, Johns AL, Patch AM, Gingras MC, et al. Genomic Analyses Identify Molecular Subtypes of Pancreatic Cancer. Nature (2016) 531(7592):47–52. doi: 10.1038/nature16965.

41. Oh K, Yoo YJ, Torre-Healy LA, Rao M, Fassler D, Wang P, et al. Coordinated Single-Cell Tumor Microenvironment Dynamics Reinforce Pancreatic Cancer Subtype. Nat Commun (2023) 14(1):5226. Epub 20230826. doi: 10.1038/s41467-023-40895-6.

42. Ma L, Tian X, Guo H, Zhang Z, Du C, Wang F, et al. Long Noncoding Rna H19 Derived Mir-675 Regulates Cell Proliferation by Down-Regulating E2f-1 in Human Pancreatic Ductal Adenocarcinoma. J Cancer (2018) 9(2):389–99. Epub 20180101. doi: 10.7150/jca.21347.

43. Zhu X, Shi C, Peng Y, Yin L, Tu M, Chen Q, et al. Thymidine Kinase 1 Silencing Retards Proliferative Activity of Pancreatic Cancer Cell Via E2f1-Tk1-P21 Axis. Cell Prolif (2018) 51(3):e12428. Epub 20171220. doi: 10.1111/cpr.12428.

44. Lammens T, Li J, Leone G, De Veylder L. Atypical E2fs: New Players in the E2f Transcription Factor Family. Trends Cell Biol (2009) 19(3):111–8. Epub 20090207. doi: 10.1016/j.tcb.2009.01.002.

45. Thurlings I, Martinez-Lopez LM, Westendorp B, Zijp M, Kuiper R, Tooten P, et al. Synergistic Functions of E2f7 and E2f8 Are Critical to Suppress Stress-Induced Skin Cancer. Oncogene (2017) 36(6):829–39. Epub 20160725. doi: 10.1038/onc.2016.251.

46. Lee S, Park YR, Kim SH, Park EJ, Kang MJ, So I, et al. Geraniol Suppresses Prostate Cancer Growth through Down-Regulation of E2f8. Cancer Med (2016) 5(10):2899–908. Epub 20160928. doi: 10.1002/cam4.864.

47. Ye L, Guo L, He Z, Wang X, Lin C, Zhang X, et al. Upregulation of E2f8 Promotes Cell Proliferation and Tumorigenicity in Breast Cancer by Modulating G1/S Phase Transition. Oncotarget (2016) 7(17):23757–71. doi: 10.18632/oncotarget.8121.

48. Ginsberg D. E2f1 Pathways to Apoptosis. FEBS Letters (2002) 529(1):122–5. doi: 10.1016/S0014-5793(02)03270-2.

49. Pützer BM, Engelmann D. E2f1 Apoptosis Counterattacked: Evil Strikes Back. Trends in Molecular Medicine (2013) 19(2):89–98. doi: 10.1016/J.MOLMED.2012.10.009.

50. Morgunova E, Yin Y, Jolma A, Dave K, Schmierer B, Popov A, et al. Structural Insights into the DNA-Binding Specificity of E2f Family Transcription Factors. Nat Commun (2015) 6:10050. Epub 20151203. doi: 10.1038/ncomms10050.

51. Lopez-Mejia IC, Fajas L. Cell Cycle Regulation of Mitochondrial Function. Curr Opin Cell Biol (2015) 33:19–25. Epub 20141112. doi: 10.1016/j.ceb.2014.10.006.

52. Benevolenskaya EV, Frolov MV. Emerging Links between E2f Control and Mitochondrial Function. Cancer Res (2015) 75(4):619–23. Epub 20150129. doi: 10.1158/0008-5472.CAN-14-2173.

53. Denechaud PD, Fajas L, Giralt A. E2f1, a Novel Regulator of Metabolism. Front Endocrinol (Lausanne) (2017) 8:311. Epub 20171110. doi: 10.3389/fendo.2017.00311.

54. Criscuolo D, Avolio R, Matassa DS, Esposito F. Targeting Mitochondrial Protein Expression as a Future Approach for Cancer Therapy. Front Oncol (2021) 11:797265. Epub 20211123. doi: 10.3389/fonc.2021.797265.

55. Liu N, Funasaka K, Obayashi T, Miyahara R, Hirooka Y, Goto H, et al. Atad2 Is Associated with Malignant Characteristics of Pancreatic Cancer Cells. Oncol Lett (2019) 17(3):3489–94. Epub 20190123. doi: 10.3892/ol.2019.9960.

56. Albahde MAH, Zhang P, Zhang Q, Li G, Wang W. Upregulated Expression of Tuba1c Predicts Poor Prognosis and Promotes Oncogenesis in Pancreatic Ductal Adenocarcinoma Via Regulating the Cell Cycle. Front Oncol (2020) 10:49. Epub 20200214. doi: 10.3389/fonc.2020.00049.

57. Kandimalla R, Tomihara H, Banwait JK, Yamamura K, Singh G, Baba H, et al. A 15-Gene Immune, Stromal, and Proliferation Gene Signature That Significantly Associates with Poor Survival in Patients with Pancreatic Ductal Adenocarcinoma. Clin Cancer Res (2020) 26(14):3641–8. Epub 20200331. doi: 10.1158/1078-0432.CCR-19-4044.

58. Meng QC, Wang HC, Song ZL, Shan ZZ, Yuan Z, Zheng Q, et al. Overexpression of Ndc80 Is Correlated with Prognosis of Pancreatic Cancer and Regulates Cell Proliferation. Am J Cancer Res (2015) 5(5):1730–40. Epub 20150415.

59. Peng YP, Zhu Y, Yin LD, Zhang JJ, Guo S, Fu Y, et al. The Expression and Prognostic Roles of Mcms in Pancreatic Cancer. PLoS One (2016) 11(10):e0164150. Epub 20161003. doi: 10.1371/journal.pone.0164150.

60. Smith SJ, Li CM, Lingeman RG, Hickey RJ, Liu Y, Malkas LH, et al. Molecular Targeting of Cancer-Associated Pcna Interactions in Pancreatic Ductal Adenocarcinoma Using a Cell-Penetrating Peptide. Mol Ther Oncolytics (2020) 17:250–6. Epub 20200408. doi: 10.1016/j.omto.2020.03.025.

61. Wijnen R, Pecoraro C, Carbone D, Fiuji H, Avan A, Peters GJ, et al. Cyclin Dependent Kinase-1 (Cdk-1) Inhibition as a Novel Therapeutic Strategy against Pancreatic Ductal Adenocarcinoma (Pdac). Cancers (Basel) (2021) 13(17). Epub 20210830. doi: 10.3390/cancers13174389.

