## Supplementary material for "E2F transcription factors promote tumorigenicity in pancreatic ductal adenocarcinoma": Legends for supplemental files

### Supplementary figure legends

**Supplementary Figure 1.** **Immunohistochemical staining patterns of E2F1 and E2F8 in PDAC**. Representative images of E2F1 expression in PDAC: A) negative E2F1 expression, B) weak positivity, C) moderate positivity, and D) strong positivity. E2F8 expression E) negative E2F8 expression, F) weak positivity, G) moderate positivity, and H) strong positivity. Original magnification: 200x.

**Supplementary Figure 2.** **Unbiased identification of cell types from an 8-week PC embryo**. Each colour represents a cell cluster from scRNA sequencing revealed by unsupervised clustering and projected on a 2-D tSNE map. Clusters have been linked to cell types using their transcriptomic signatures

**Supplementary Figure 3. Identification of cell types from PDAC tissues**. Localization of E2F1^+^ (A), E2F8^+^ (B) and E2F1^+^E2F8^+^ (C) cells for each cell type clusters.

**Supplementary Figure 4. Transfection efficiency determined by RT-qPCR analyses.** E2F1 (A) and E2F8 (B) knockdown efficiency 48h or 7 days after transfection. Expression values were calculated applying the −2∆∆CT algorithm. Estimated relative quantities were normalized for the expression value of the endogenous genes GAPDH, S18 and TBP and calibrated to the negative control samples.

### Supplementary table legends

**Table S1.** Primer sequences for qPCR.

**Table S2.** Differential expression analysis of E2F genes in adult and fetal pancreas.

**Table S3.** Overall survival analyses

**Table S4.** E2F1 co-expression analyses in embryo tissues using Spearman's rank correlation method.

**Table S5.** E2F8 co-expression analyses in embryo tissues using Spearman's rank correlation method.

**Table S6.** E2F1 co-expression analyses in normal adult tissues using Spearman's rank correlation method.

**Table S7.** E2F1 co-expression analyses in PDAC tissues using Spearman's rank correlation method.

**Table S8.** E2F8 co-expression analyses in PDAC tissues using Spearman's rank correlation method.

**Table S9.** Genes found in E2F1 ChiP-seq dataset and co-expressed with E2F1 in PDAC, organized by pathways and classified by significance.

**Table S10.** Genes found in E2F8 ChiP-seq dataset and co-expressed with E2F8 in PDAC, organized by pathways and classified by significance.

**Table S11.** References listing the role during carcinogenesis of the most significant genes identified in table S8.

**Table S12.** References listing the role during carcinogenesis of the most significant genes identified in table S9.
